# Structure-guided targeting of the GATAD2A-CHD4 interaction within the MBD2-NuRD complex results in high levels of HbF in adult erythroid cells

**DOI:** 10.64898/2026.07.17.739209

**Authors:** Shengzhe Shang, Sangam Goswami, Torry Li, He Wang, Christopher Travis, Mikhail Dozmorov, David C Williams, Gordon D Ginder

## Abstract

Fetal hemoglobin (HbF) expression is silenced postnatally in adult erythroid cells. Sufficiently increased expression of HbF has been shown to overcome the pathophysiologic sequelae of both sickle cell disease and beta-thalassemia. As the MBD2a-NuRD chromatin remodeling complex is required for silencing of HbF, the present studies were aimed at exploring a potential therapeutic approach for disrupting this complex. AlphaFold 3 and a recent crystal structure were employed to predict the critical interaction domains linking GATAD2A in the histone deacetylase core subcomplex (HDCC) of NuRD and the CHD4 ATPase which has been shown to be required for silencing of the fetal gamma-globin (*HBG*) genes. The two predicted critical domains, the CR2 helical domain of GATAD2A and the C-terminal domains 1 and 2 (C1b and C2ab) of CHD4, were validated by *in vitro* biophysical studies. Mutation of two amino acids in the CR2 helical domain of the endogenous GATAD2A gene in HUDEP-2 cells resulted in dissociation of CHD4, loss of repressive chromatin over the HBG promoter and ~40% HbF levels compared to < 1% in control cells. Strikingly, enforced expression of a peptide containing the helical portion of the CR2 domain of GATAD2A in both HUDEP-2 cells and primary adult erythroid cells resulted in high levels of HbF, with up to ~75% HbF compared to mutant peptide control level of ~9% in the latter without perturbing erythroid differentiation. These results suggest that targeting the critical interaction domains of GATAD2A and CHD4 with a macrocyclic peptide or small molecule may lead to much needed small molecule therapeutics for sickle cell disease.

**Key Points:** Association of CHD4 with the HDCC core of the MBD2-NuRD chromatin remodeling complex is required for silencing of HbF expression in adult human erythroid cells

Genetic alteration or enforced peptide expression of a critical helical domain of GATAD2A results in dissociation of CHD4 from the MBD2-NuRD complex and high-level expression of HbF.

## Introduction

The regulation of hemoglobin expression during vertebrate ontogeny has served as a leading model of developmental gene regulation. In humans, the predominant beta-like globin chain and its corresponding hemoglobin type undergo a sequential switch from embryonic epsilon (Hb Gower) to fetal gamma (HbF) to predominantly beta (HbA) with a minor component of delta (HbA2) postnatally. It is well documented that persistent expression of HbF postnatally can substitute for decreased levels of HbA, as occurs naturally in hereditary persistence of fetal hemoglobin (HPFH). Similarly, it has been shown that sufficient levels of HbF can overcome the molecular consequences of the defective mutant beta chain in HbS in sickle cell disease (SCD) as well as the severe deficit of beta globin in biallelic β-thalassemia.^1,2^ These observations have inspired research to understand and potentially reverse the silencing of HbF therapeutically in these common single-gene disorders.

Several major repressive transcription factors necessary for γ-globin gene (HBG) and thus HbF silencing have been characterized, including two of the most potent, BCL11A ^3–5^ and ZBTB7A,^6^ also known as leukemia-related factor (LRF), among many others that have been identified.^7^ In addition to transcription factors, epigenetic factors, including DNA methylation at the *HBG* promoters,^8–10^ MBD2a and its associated NuRD chromatin remodeling complex,^11–15^ WIZ,^16^ G9a,^17,18^ PRMT5,^14,19^ LSD1,^20^ and histone deacetylase,^21,22^ have been shown to mediate HbF silencing. Although a large number of factors have been shown to be involved in HbF silencing^7,23^, a central mechanism appears to be mediated through a multi-protein complex anchored at the proximal *HBG* promoter by the co-dependent combination of binding of BCL11A and ZBTB7A, CpG methylation, and methylation-dependent binding of the MBD2a-NuRD chromatin remodeling complex at the promoter.^9^

Despite the success of gene therapy aimed at depleting BCL11A levels at the HBG promoters to therapeutically increase HbF in patients with SCD,^24–27^ there are economic and logistical challenges to the widespread adoption of this modality. As such, there is a pressing need for more effective small-molecule therapeutics than the current standard, hydroxyurea (HU), to reach the millions of SCD patients worldwide, particularly in resource-limited regions where the majority of these individuals reside.

Based on the absolute requirement of MBD2a-NuRD for *HBG* silencing^11,13–15^ and the relatively mild phenotype of MBD2 knockout mice,^28^ MBD2a-NuRD presents a potentially attractive target for small-molecule therapeutics. Previous pre-clinical studies have shown that depletion of key components of MBD2a-NuRD, and in particular the CHD4 ATPase component, results in 40-50% HbF levels in adult primary erythroid cells as well as in the HUDEP-2 human adult erythroid cell model.^12,13,15,29–32^

A challenge in disrupting the MBD2a-NuRD complex is the fact that it requires inhibition of protein-protein interactions, which have often been considered undruggable despite examples in which this barrier has been overcome.^33,34^ To discover proof-of-principle agents capable of relieving silencing by disrupting this complex, we have carried out extensive structure-function studies to identify potential targets for disruption.^12,13,35^

In this report, we describe a combination of structural and biophysical validation experiments used to characterize a critical interaction between the CR-2 domain of GATAD2A and the C-terminal domains (C1b and C2ab) of CHD4 that tether the latter to the MBD2a-NuRD complex.^36^ Functional biochemical and genetic assays validated the key role of this protein-protein interface and demonstrated that mutation of critical amino acids in the helical portion of the CR-2 domain of GATAD2A, or expression of an interfering peptide containing this domain, can disrupt its interaction with CHD4 and induce high levels of HbF in both HUDEP-2 cells and primary CD34^+^ HSPC (Hematopoietic Stem and Progenitor Cell) derived adult erythroid cells without perturbing differentiation or cell viability.

## Methods

### Protein Expression and Purification

#### GATAD2A-CR2

We cloned DNA encoding the internal α-helix and Zn finger domain from the GATAD2A-CR2 region (residues 371-444) into the pET28a vector with N-terminal hexahistidine and maltose binding protein fusions and a C-terminal TwinStrep tag. For expression of the isolated CR2-helix, we cloned DNA encoding GATAD2A residues 371-404 into the pET32a(+) vector with N-terminal thioredoxin and hexahistidine tags and a C-terminal TwinStrep tag. Both constructs include a tobacco etch virus (TEV) protease site (ENLYFQG) immediately preceding the expressed domain, such that TEV protease digestion removes the N-terminal tags. We transformed the vectors into Rosetta2(DE3), grew the bacteria under ampicillin selection at 37 °C until an optical density at 600 nm of ~0.9, and induced with 1 mM isopropyl β-d-1-thiogalactopyranoside (IPTG) for ~2-3 hours.

For the isolated CR2-helix, we lysed the bacteria in 30 mL of 20 mM Tris, pH 8.0, 1 M NaCl, 1 mM DTT, with sonication, and purified the protein by nickel affinity chromatography, eluting with 20 mM Tris, pH 8.0, 1 M NaCl, 300 mM imidazole, and 1 mM DTT. The N-terminal tags were removed by overnight digestion with TEV protease at room temperature, followed by purification over a Streptactin XT 4 Flow column (IBA Lifesciences), and finally size-exclusion chromatography (Superdex 75 Increase 10/300, Cytiva) in 20 mM Tris, pH 8.0, 150 mM NaCl, 1 mM DTT.

For the CR2 region, we followed a similar strategy, except that the initial step was purification on an MBPTrap column (Cytiva) rather than nickel affinity chromatography. The protein was eluted with 10 mM maltose in lysis buffer, and the subsequent purification was performed as above.

#### CHD4-C1bC2a

We cloned the C1b (residues 1395-1519) and C2a (residues 1693-1810) domains of human CHD4 protein, connected with a four amino acid (GGGS) linker, into the same modified pET32a (+) vector as for GATAD2A-CR2helix, which includes N-terminal thioredoxin and hexahistidine tags with a TEV protease site. We transformed the vector into Rosetta2(DE3), grew the bacteria to an optical density at 600 nm of ~0.6, lowered the temperature to 20 °C, and induced protein production with 0.4 mM IPTG overnight (16-20 hours). We lysed the bacteria by sonication in 30 mL of buffer A (20 mM Tris, pH 8.0, 0.5 M NaCl, 1 mM dithiothreitol (DTT), 5% glycerol), plus 1 mM PMSF, and purified by nickel affinity chromatography. The N-terminal tags were removed by TEV protease digestion while dialyzing against buffer A overnight, followed by a second nickel affinity chromatography step, and a final purification by size exclusion chromatography (Superdex 75 Increase 10/300, Cytiva) in buffer A.

#### Ternary Complex (CHD4-C1bC2ab + GATAD2A-CR2 + CDK2AP1)

We purified the ternary complex as described previously.^37^ Briefly, we subcloned the full CR2 domain from GATAD2A (with an N-terminal TwinStrep tag and TEV protease site) and the C1bC2ab domains of CHD4 into the pETDuet-1 vector. We subcloned CDK2AP into the pCDF vector (with N-terminal thioredoxin and hexahistidine tags and a TEV protease site). We co-transformed the vectors into Rosetta2(DE3), grew until an optical density at 600 nm of ~0.6, lowered the temperature to 20 °C, and induced with 0.4 mM overnight (16-20 hours). We lysed by sonication in 30 mL of buffer A (as above) and purified by sequential affinity chromatography over nickel (5 mL HisTRAP, Cytiva) and streptactin (5 mL Streptactin XT 4Flow, IBA Lifesciences) resins. After overnight TEV protease cleavage at room temperature, we purified the complex by nickel affinity and size-exclusion chromatography in buffer A.

### Isothermal Titration Calorimetry

The purified GATAD2A-CR2 domains (50-100 μM) were titrated (20-27 injections, 2 μL per injection, at 30 °C) into CHD4-C1bC2a (5-10 μM) in buffer A, and the heat was measured on a PEAQ-ITC calorimeter (Malvern). The resulting isotherms were fit to a 1:1 binding model with the manufacturer’s software.

### Differential Scanning Fluorimetry (nanoDSF)

We mixed CHD4-C1bC2a or the ternary complex (10 μM) with wild-type and mutant GATAD2A-CR2helix (20 μM or 100 μM, respectively), and differential scanning fluorimetry was recorded on a Prometheus NT.48 (NanoTemper). The samples were heated from 20 to 95 °C (1 °C per minute), and tryptophan fluorescence was measured at 350 nm and 330 nm. We identified the melting temperature by taking the first derivative of the fluorescence intensity ratio (350 nm/330 nm) and fitting the peak maxima using an in-house Python script (https://github.com/dcwilljr/plotNanoDSF).

### NanoBRET assays

We cloned the CHD4 C1bC2ab domains and GATAD2A CR2 domains into the pNLF1-C and pHTC vectors (Promega) in frame with C-terminal NanoLuciferase and HaloTag fusions, respectively (**Figure 1D**). We measured intracellular bioluminescent energy transfer following the manufacturer’s protocol, as previously described.^35^

**Figure 1.**
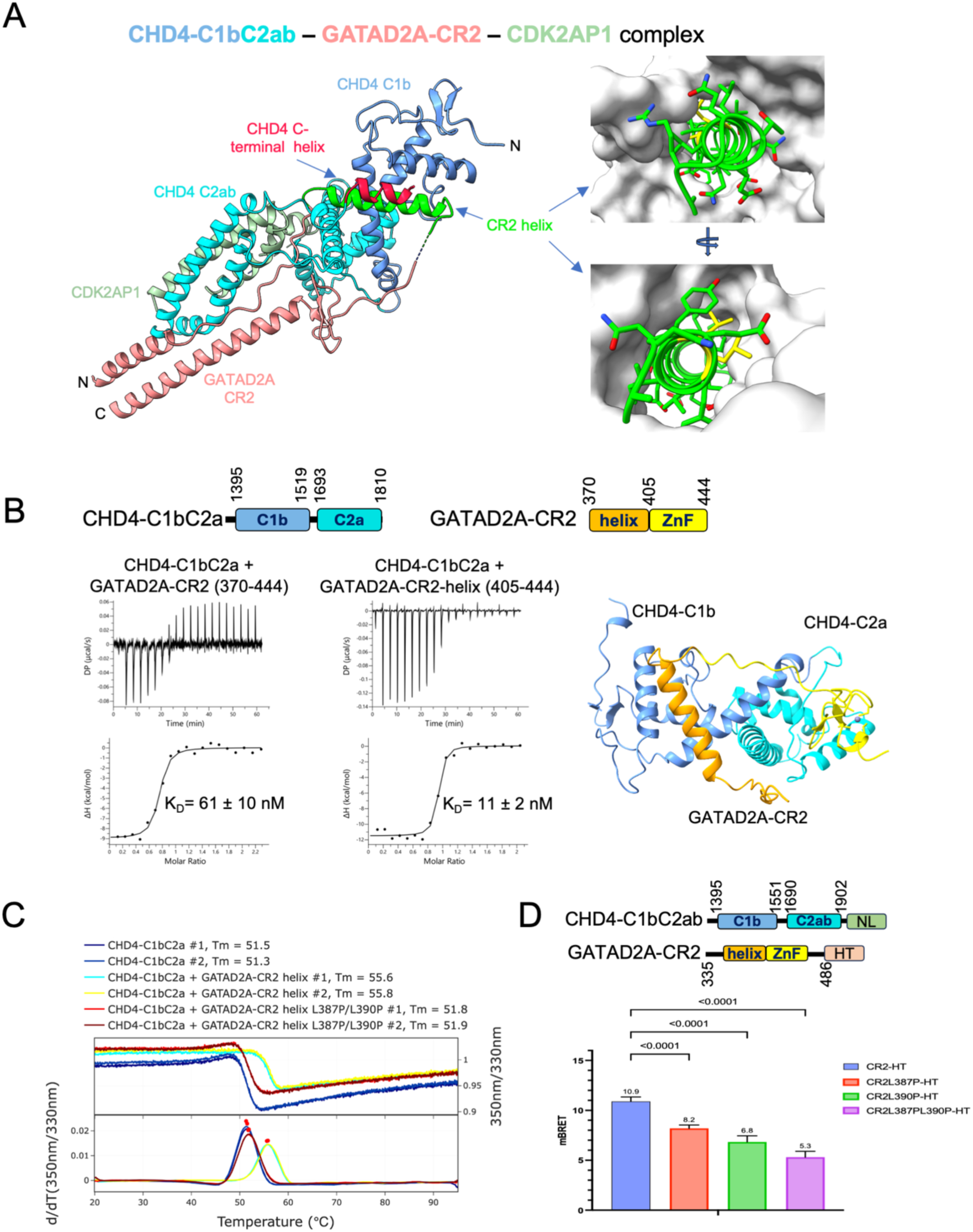
The GATAD2A CR2 region binds tightly to the C-terminal domains of CHD4 through a short helix – protein interaction. A. A ribbon diagram (left) of the crystal structure of the ternary complex shows the interaction between GATAD2A-CR2 region (light coral) and CHD4-C1b (cornflower blue) and CHD4-C2ab (cyan) domains and CDK2AP1 (light green). The GATAD2A-CR2 helix (green) binds at the interface between the CHD4 C1b and C2a domains. The CHD4 C-terminal helix (red) binds atop the CR2 helix, such that CHD4 completely surrounds it, as shown in the right panels. B. Isothermal titration calorimetry binding isotherms show that the GATAD2-CR2 region encompassing both the CR2-helix (orange) and Zn-finger (yellow), and the isolated GATAD2-CR2 helix bind to CHD4-C1bC2a domains with high affinity (n = 2). Molecular diagrams (above) and a molecular model (right) depict the constructs used for ITC measurements. C. NanoDSF measurements of CHD4-C1bC2a in isolation and bound to wild-type and mutant (L387P/L390P) GATAD2A-CR2 helix show that the wild-type helix shifts the melting temperature by 4.3 °C, whereas the mutant helix shifts it by only 0.5 °C. D. NanoBRET measurements reveal a large signal consistent with an intracellular interaction between CHD4-C1bC2ab and wild-type GATAD2A-CR2, while L387P and L390P significantly reduce the signal, with the double mutation showing the largest reduction.

### Cell culture and differentiation, Western blot, IP, ChIP-qPCR, NOMe-seq

HUDEP-2 cell culture and differentiation methods, as well as lentivirus packaging and transduction, Western blot, immunoprecipitation, ChIP-qPCR, NOMe-seq, have been described previously ^14^.

### Antibodies and reagents

Antibodies used in this study are listed in SI Appendix, Table S1. Protein G Dynabeads were from Thermo Fisher. Protein A/G agarose beads were from Santa Cruz Biotechnology. StemSpan and SFEM II were purchased from StemCell Technologies. Recombinant human stem cell factor (SCF) and Doxycycline (DOX), Dexamethasone (DEX), recombinant human insulin, heparin and human AB serum were obtained from Sigma Aldrich. Recombinant Erythropoietin (EPO) was purchased from Virginia Commonwealth University (VCU) main hospital. Holo-human transferrin (HTF) was purchased from Prospec.

### Plasmids

Prime editor system was purchased from Addgene (pCMV-PEmax, #174820. pU6-tevopreq1-GG-acceptor, # 174038). Packaging plasmid psPAX2 (Addgene #12260) and pMD2.G (Addgene #12259), lentiviral pLV203 and CRISPR plasmid LentiCRISPRv2 were as previously saved in our lab (Addgene #98290).

### Lentiviral-mediated “Add-back” of GATAD2A wild type and mutants in GATAD2A null cells

Lentivirus containing wildtype GATAD2A or GATAD2A mutants infected in GATAD2A knockout HUDEP-2 cells. The assessment of exogenous GATAD2A expression in HUDEP-2 GATAD2A KO cells was carried out by Western immunoblotting five days post lentiviral infection as described ^12^.

### Prime editing of GATAD2A gene in HUDEP-2

The lentiviral prime editor and epegRNA targeting GATAD2 residues L387 and L390 were constructed from original plasmids obtained from Addgene. The prime editing system was then introduced into HUDEP-2 cells. Transduced cells were selected with puromycin for 3 days post-infection. Genomic DNA was extracted to assess editing efficiency by PCR and Sanger sequencing. Puromycin-selected cells were seeded into 96-well plates for single-cell cloning. Individual clones were screened by PCR and Sanger sequencing to determine editing outcomes. Both edited and unedited clones were differentiated in parallel for 6 days, after which mRNA was collected for qPCR analysis to quantify globin gene expression.

### RNA-Seq and data analysis

CD34^+^ hematopoietic stem and progenitor cells (HSPCs) were transduced with lentivirus expressing either wild-type (WT) or mutant GATAD2A CR2 during the expansion phase. Following transduction, cells were maintained in expansion medium until day 6 as previously described.^14^ Cells were then induced to undergo erythroid differentiation using the previously described 8-day differentiation protocol.^38^ Briefly, CD34^+^ HSPCs were cultured at a density of 0.5–1 × 10^6^ cells/mL in erythroid differentiation medium (EDM) consisting of IMDM (Thermo Fisher Scientific) supplemented with 3% human AB serum, 2% fetal bovine serum (FBS), 1% GlutaMAX, 1% penicillin-streptomycin, 10 ng/mL recombinant human stem cell factor (SCF), 200 μg/mL human holo-transferrin (HTF), 1 ng/mL human interleukin-3 (IL-3), 3 IU/mL recombinant human erythropoietin (EPO), 10 μg/mL insulin, and 1 μM dexamethasone (DEX). After 3 days of differentiation, cells were harvested, and total RNA was extracted using the Total RNA Mini Prep Kit (New England Biolabs). The purified RNA was subsequently used for RNA sequencing (RNA-seq). The sequencing was done by Novogene with NovaSeq X Plus Series (PE150), 6G raw data per sample (three replicates for CR2 WT and three replicates for CR2 mutant). RNA-seq data was processed using Nextflow rnaseq pipeline (v.3.12) which performs QC, adapter trimming, alignment to reference genome GRCh38 with contigs and “sponge” sequences using STAR and transcript quantification using RSEM. Differential gene expression analysis was conducted using Bioconductor package edgeR v4.2.1, based on normalized and filtered gene counts. Genes exhibiting statistically significant changes (adjusted p-value < 0.05) were further analyzed using downstream bioinformatics approaches.

## Results

### The CR-2 domain of GATAD2A binds to the C1bC2a domains of CHD4

The NuRD complex consists of seven core components that form two distinct functional sub-complexes: the histone deacetylase core complex (MTA1/2/3, HDAC1/2, RBBP4/7) and chromatin remodeling complex (CHD4, CDK2AP1) that are bridged by the MBD2/3 and GATAD2A/B proteins. Previous work has shown that the CR-2 domain of GATAD2A binds to the C-terminal region of CHD4.^36^

We sought to determine whether disrupting this interaction would relieve NuRD-dependent silencing of fetal hemoglobin expression. We first measured the binding between CHD4-C1bC2a and GATAD2A-CR2 by isothermal titration calorimetry (ITC). As shown in Figure 1B, a peptide containing the GATAD2ACR2 Zn-finger and central helix binds to CHD4-C1bC2a with high-affinity (K_D_ = 61 ± 10 nM), whereas the CR2 helix in isolation binds to CHD4-C1bC2a with an approximately 5.5-fold increase in affinity (K_D_ = 11 ± 2 nM). We mutated two leucine residues to proline (L387P and L390P) in the CR2 helix to disrupt helix formation. We measured the change in thermal stability upon binding for the native and mutant CR2 helix by differential scanning fluorometry (NanoDSF, Nanotemper Prometheus NT.48). The CHD4-C1bC2a domain shows a melting temperature of 51.4 ± 0.1 °C in isolation, 55.7 ± 0.1 °C bound to GATAD2A-CR2 helix (ΔTm = 4.3 °C), and 51.9 ± 0.1 °C bound to GATAD2A-CR2 helix L387P/L390P (ΔTm = 0.5 °C) (Figure 1C). We then tested whether the proline mutations disrupted binding between the full CHD4-C1bC2ab domain, which includes the C-terminal helix that binds to the CR2-helix, and the full GATAD2A-CR2 region using an in-cell bioluminescent energy transfer assay (NanoBRET). Co-expressing the CHD4-C1bC2ab with a C-terminal NanoLuc and GATAD2A-CR2 with a C-terminal HaloTag fusion yields a strong BRET signal. The GATAD2A-CR2 L387P and L390P mutations significantly reduce the BRET signal, with the combined mutations leading to the largest reduction.

### Disrupting the interaction between the CR2-helix and the CHD4-C1bC2ab reduces the stability of the ternary complex at the site of interaction

In recent related studies, we determined the structure of a complex between the C-terminal domain of CHD4, the CR2 domain of GATAD2A, and the CDK2AP1 protein.^37^ The structure of this complex shows that a small helix from GATAD2A-CR2 (amino acids 370-405) makes extensive contacts at the interface between the C1b and C2a domains of CHD4 (Figure 1A). In addition, we identified macrocyclic peptides that competitively inhibit binding of the CR2-helix to CHD4.

For this report, we used bacterial expression constructs to co-express and purify a ternary complex comprising CHD4-C1bC2ab, GATAD2A-CR2, and CDK2AP1.^37^ To determine whether the CR2-helix could destabilize the formation of the ternary complex, we measured the melting temperature of the complex in isolation and in the presence of wild-type and mutant GATAD2ACR2-helix peptides (Figure 2A). The results show that adding excess wild-type peptide reduces the melting temperature from 56.7 ± 0.1 °C to 53.1 ± 0.1 °C (ΔTm = −3.6 °C). In contrast, the mutant peptide causes slight broadening of the transition without a large reduction in melting temperature (56.3 ± 0.1 °C, ΔTm = −0.4 °C). These results indicate that the isolated GATAD2A-CR2 helix can disrupt formation of the NuRD complex.

**Figure 2.**
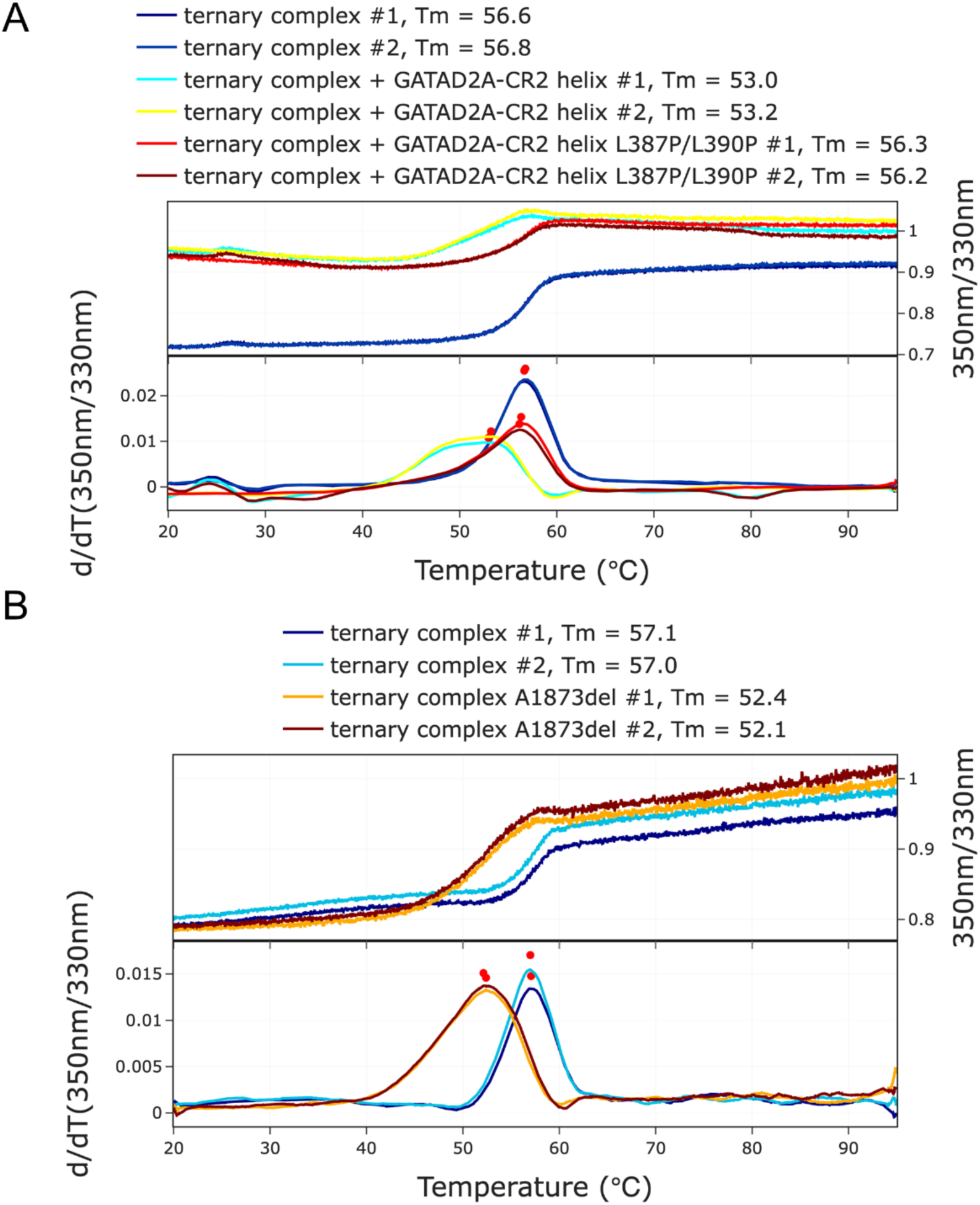
Disrupting the interaction between GATAD2A-CR2 helix and CHD4-C1bC2ab destabilizes ternary complex formation. A. NanoDSF measurements of the ternary complex show that the addition of the wild-type GATAD2A-CR2 helix broadens the melting curve and reduces the observed melting temperature by 3.6 °C, whereas the mutant helix shifts it by only 0.5 °C. B. NanoDSF measurements show that the CHD4 A1873 deletion mutation destabilized the ternary complex, reducing the observed melting temperature by 4.8 °C.

Previous work by the Bauer group identified deletion mutants at sites immediately preceding the CHD4 C-terminal helix that increase fetal hemoglobin expression without decreasing cell fitness, in both HUDEP-2 cells and transgenic mice. We reasoned that these mutants destabilize the interaction between the C-terminal helix and GATAD2A-CR2 helix (Figure 1A). To test this hypothesis, we introduced the A1873del mutation into the CHD4-C1bC2ab expression construct and purified the mutant ternary complex. As shown in Figure 2B, the deletion mutant shows a large reduction in thermal stability (ΔTm = −4.8 °C). Together, these studies show that the interaction between the GATAD2A-CR2 helix and CHD4-C1bC2ab represents a structural vulnerability that could be targeted to reduce NuRD complex stability and induce the expression of fetal hemoglobin.

### Mutation of two amino acids in the CR-2 domain helix of GATAD2A results in its dissociation of CHD4 from the MBD2-NuRD histone deacetylase core subcomplex and relieves *HBG* silencing

To confirm the functional importance of the structural predictions regarding the interaction between the CR-2 helical domain of GATAD2A and the C-terminal domains 1 and 2 of CHD4 (CHD4 CTD 1/2), two independent approaches were taken. In the first of these *GATAD2A* was knocked out in HUDEP-2 cells followed by enforced expression of either wild type (WT) or mutated *GATAD2A* bearing 2 proline amino acid substitutions at positions L387 and L390 in the CR-2 helical domain. As shown in Figure 3A and 3B, only add back of WT GATAD2A resulted in partial restored silencing of *HBG* expression. Given the limited ability of enforced expression add back experiments to show the full magnitude of the effects of protein mutations,^13^ prime editing^39^ was carried out to introduce the same two proline substitutions in the endogenous *GATAD2A* gene at positions L387 and L390 (Supplemental Figure 1A). As shown in Figure 3C, these mutations resulted in over a 140-fold increase in *HBG* mRNA expression and ~40% *HBG/HBG+HBB* mRNA levels (Figure 3D) compared to no significant effect in prime editor no guide control cells. The same amino acid substitutions in the endogenous GATAD2A protein resulted in a 50% decrease in *HBB* mRNA expression (Figure 3E). HPLC analysis of hemoglobin levels, shown in Figure 3F confirmed an increase in HbF level to 39% of total hemoglobin, consistent with the increase in *HBG* mRNA in the mutant HUDEP-2 cells. L387P/L390P amino acid substitutions in the CR-2 helical domain of endogenous GATAD2A did not perturb expression of the erythroid differentiation markers CD71 and CD235 (Supplemental Figure 1B). Analysis of multiple clones of HUDEP-2 cells bearing the L387P/L390P mutation on only one allele showed variable but consistently increased levels of *HBG* RNA (Supplemental Figure 1C) suggesting that haploinsufficiency of wild type GATAD2A was able to impart partial relief of *HBG* silencing. Western blots of GATAD2A immunoprecipitation pulldowns demonstrated loss of binding of GATAD2A to CHD4 in the L387P/L390P mutant HUDEP-2 cells, while not perturbing its binding to MBD2 or MTA2, which are components of the histone deacetylase core (HDCC) subcomplex of MBD2a-NuRD (Figure 3G). This result suggests that disruption of the CR-2 domain of GATAD2A does not affect the normal interaction of its CR-1 region which tethers it to MBD2 and the HDCC core subcomplex.^12^

**Figure 3.**
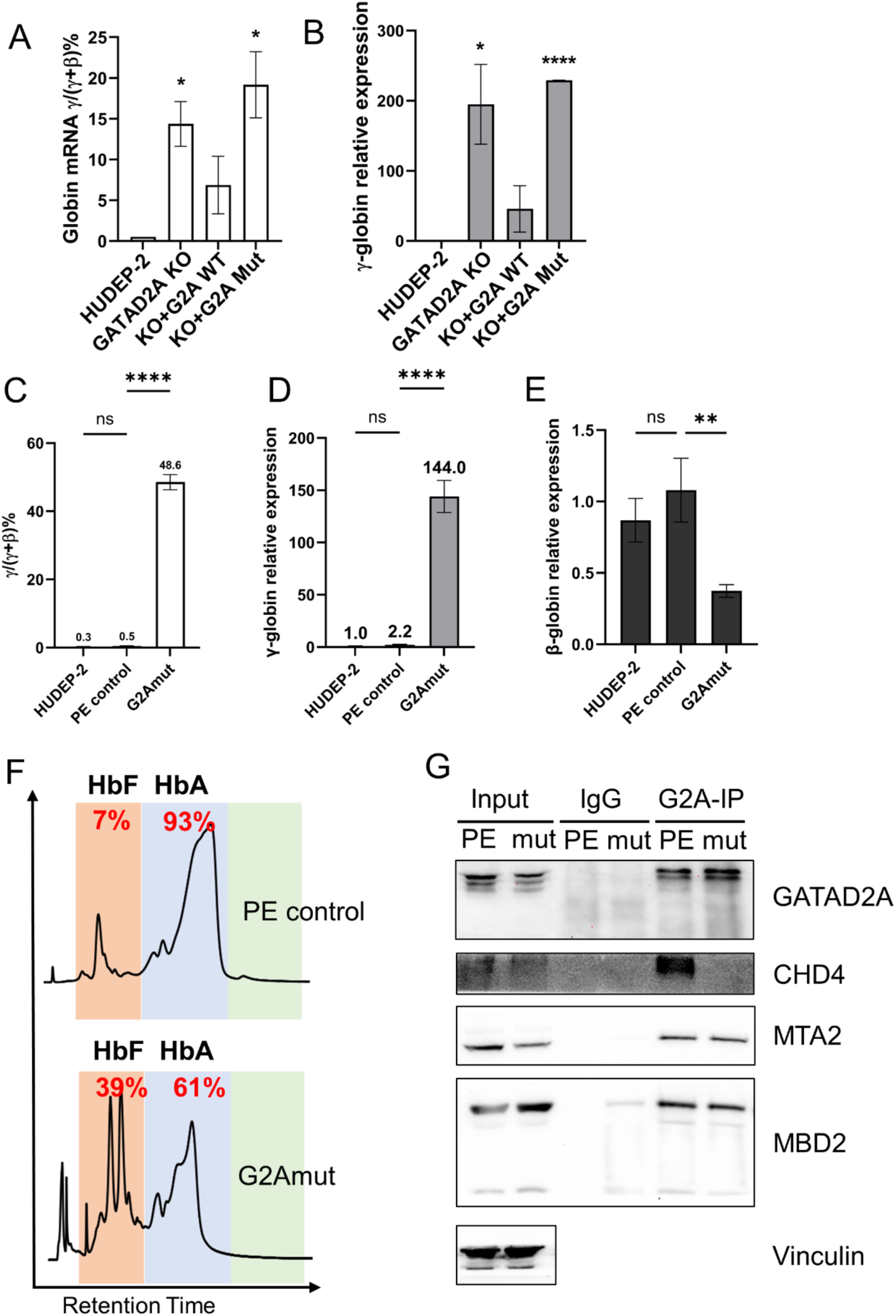
The GATAD2A L387P/L390P mutations disrupt the interaction between CHD4 and the HDCC NuRD subcomplex. A–B qPCR assay of γ/ (γ+β) mRNA ratio and relative γ-globin mRNA level in enforced expression of wild-type and double mutants GATAD2A HUDEP-2 cells. C–D. E qPCR assay of γ/ (γ+β) mRNA ratio and relative γ, β-globin mRNA level in PE edited GATAD2A mutant HUDEP-2 cells. F. HPLC analysis showing elevated HbF levels in PE-edited GATAD2A mutant HUDEP-2 cells compared with PE control HUDEP-2 cells. G. Co-IP and Western blot analyses showing that only CHD4, but not other NuRD complex components, was dissociated from the NuRD complex in GATAD2A mutant cells.

### Mutation of the CR-2 helical domain of GATAD2A results in loss of occupancy of CHD4, repressive histone marks, and repressive nucleosomes positioned over the *HBG* promoters

Chromatin immunoprecipitation Q-PCR assay showed that the L387P/L390P amino acid substitutions in CR-2 result in loss of binding of CHD4, as well as loss of the repressive histone mark H3K9me3 and acquisition of increased binding of the active chromatin marks H3K4me3 and H3K9ac at the *HBG* promoter (Figure 4A). NOMe-Seq assay showed that the disruption of the interaction between GATAD2A and CHD4 in the MBD2a-NuRD complex resulted in loss of nucleosome occupancy of the *HBG* promoter to a similar degree as observed previously in cells in which MBD2a-NuRD binding to the methylated promoter was prevented by knockout of MBD2^14^ (Figure 4B). These results confirm the important role of CHD4 in the ability of the MBD2a-NuRD complex to establish a repressive chromatin environment at the HBG promoters.

**Figure 4.**
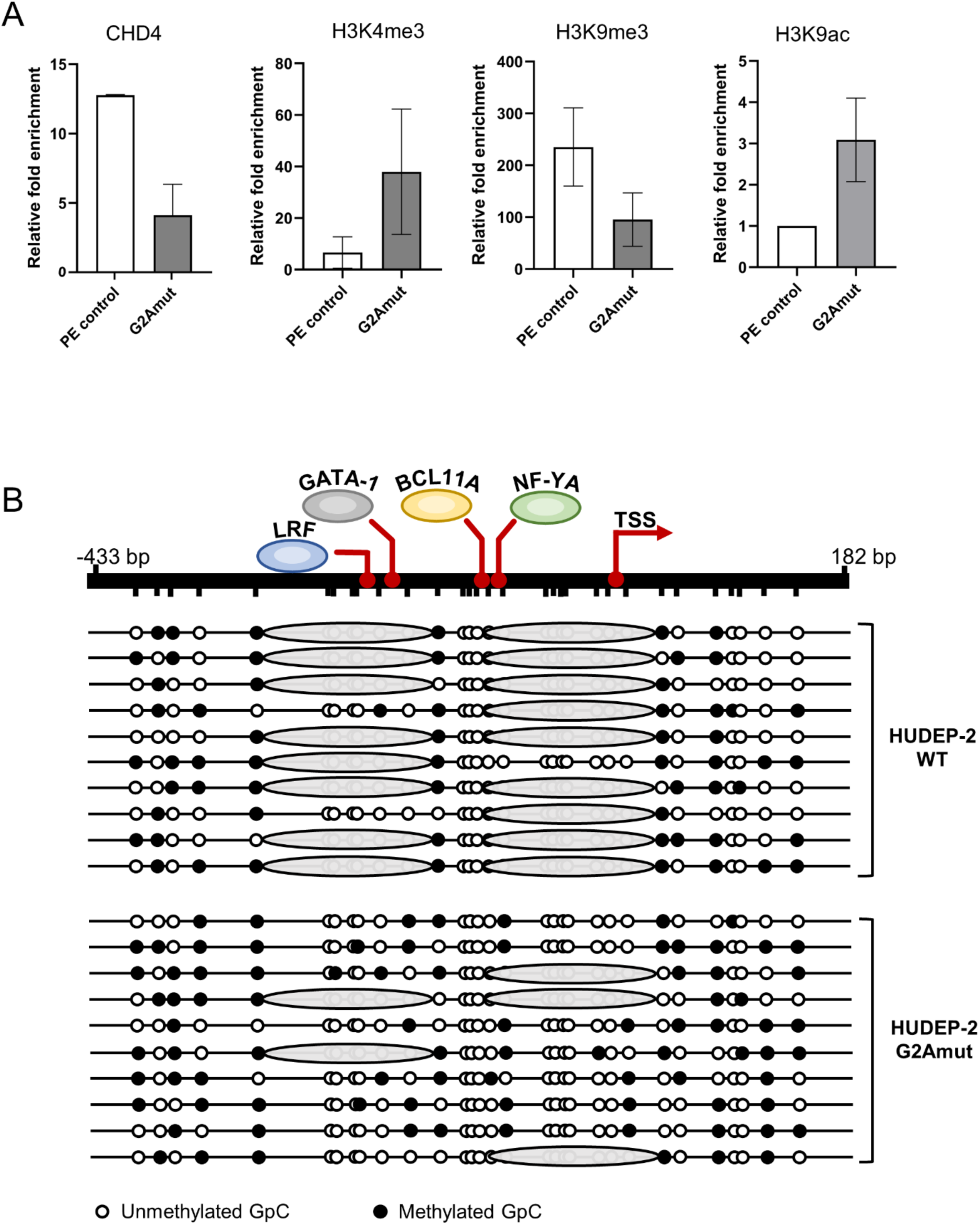
PE edited HUDEP-2 cells have decreased occupancy of CHD4, loss of repressive histone marks and loss of nucleosomes at the HBG promoter. A. ChIP-qPCR analysis showing reduced CHD4 occupancy at the HBG promoter. The activating histone marks H3K4me3 and H3K9ac were increased, whereas the repressive mark H3K9me3 was decreased at the HBG promoter consistent with an inactive chromatin environment. B. NoME-seq analysis demonstrating nucleosome loss across the HBG promoters in GATAD2A mutant HUDEP-2 cells compared to wild type cells, similar to that observed in MBD2-knockout cells.

### An interfering peptide containing the amino acids of the helix within the CR-2 domain of GATAD2A relieves *HBG* silencing without perturbing erythroid differentiation

Enforced expression in HUDEP-2 cells of a 31 amino acid peptide from the CR-2 helix of GATAD2A resulted in a 15% ratio of *HBG/HBG+HBB* mRNA and a 22-fold increase in relative *HBG* mRNA level in HUDEP-2 cells compared to 1% ratio in cells in which the CR-2 helix peptide with the L387/390P mutations as shown in Figure 5A and 5B. Western blots of strep-tagged GATAD2A CR2 peptide immunoprecipitation pulldown showed the ability of wild type CR-2 helix peptide but not L387/L390P mutant peptide to bind to CHD4 demonstrating an on-target effect of the peptide (Figure 5C).

**Figure 5.**
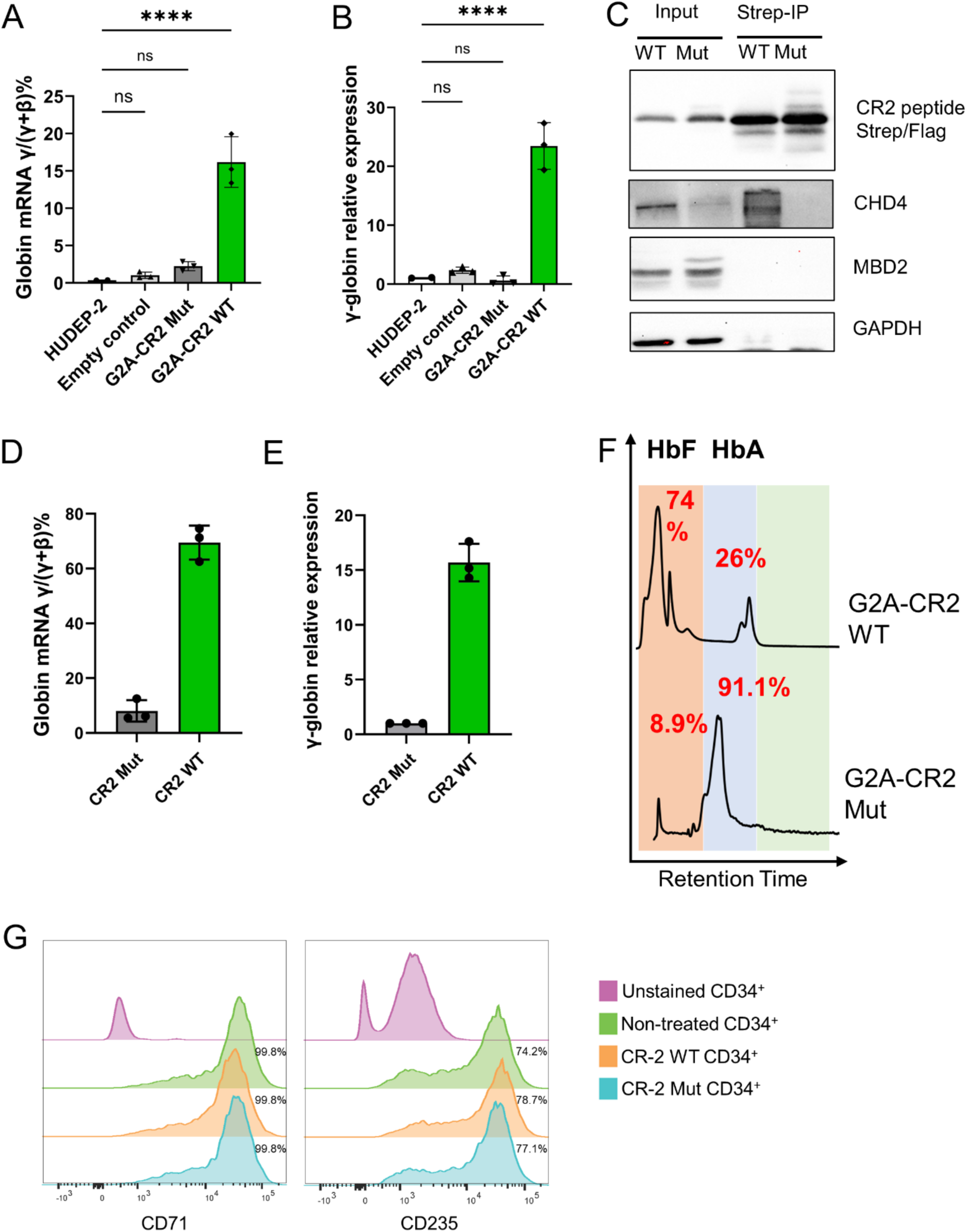
Expression of a small peptide derived from the GATAD2A CR2 helix competitively inhibits the interaction between CHD4 and the HDCC NuRD subcomplex. A–B. qPCR analysis of the γ/(γ+β) mRNA ratio and relative γ-globin mRNA levels in HUDEP-2 cells expressing wild-type or mutant GATAD2A CR2 helix peptides. C. Western blot analysis of immunoprecipitation of Strep-tagged peptides showing that only the wild-type CR2 helix peptide, but not the mutant peptide, is able to pull down CHD4. D–E. qPCR analysis of the γ/(γ+β) mRNA ratio and relative γ-globin mRNA levels in CD34+ cells expressing wild-type or mutant GATAD2A CR2 helix peptides. F. HPLC analysis demonstrating similarly elevated HbF levels in CD34+ HSPC derived erythroid cells overexpressing wild-type but not mutant CR2 helix peptide, G. Flow cytometry analysis of the erythroid differentiation markers CD71 and CD235a in CD34+ HSPC derived cells overexpressing wild-type or mutant GATAD2A CR2 helix peptides showing unperturbed differentiation.

To determine the effect of the CR-2 helix interfering peptide in primary human erythroid cells, lentiviral vector mediated enforced expression was carried out in CD34^+^ HSPCs followed by erythroid differentiation. As shown in Figure 5D and 5E, enforced expression of WT but not L387P/L390P mutant peptide resulted in ~70% *HBG/HBG+HBB* mRNA ratio and a ~15-fold increase in *HBG* mRNA, with a corresponding increase to 74% HbF (Figure 5F).

Flow cytometric analysis of differentiated HSPC-derived erythroid cells demonstrated that enforced expression of the WT CR-2 peptide did not perturb erythroid differentiation (Figure 5G)

### Enforced expression of CR-2 peptide in CD34^+^ HSPCs affects expression of only a limited number of genes after erythroid differentiation

RNA-seq analysis was carried out on the CD34^+^ HSPCs in which either the WT CR-2 helix peptide or the CR-2 mutant peptide with L387P/L390P amino acids substitutions was introduced and which displayed the corresponding *HBG/HBG+HBB* mRNA expression levels shown in Figure 5D.

As shown in the volcano plot (Figure 6A), only a limited number of genes were significantly affected. Among the most upregulated of these were *HBG1* and *HBG2* as expected from the Q-PCR data in Figure 5. Also upregulated were the embryonic β-type globin gene, *epsilon* (HBE), and the embryonic α-type globin gene, *zeta* (HBZ). The top 50 genes differentially upregulated by wild type versus mutant CR-2 peptide enforced expression are shown in the heat map (Figure 6B). There was a total of 89 genes significantly upregulated and 35 genes significantly downregulated by enforced expression of wild type CR2 peptide versus mutant CR2.

**Figure 6.**
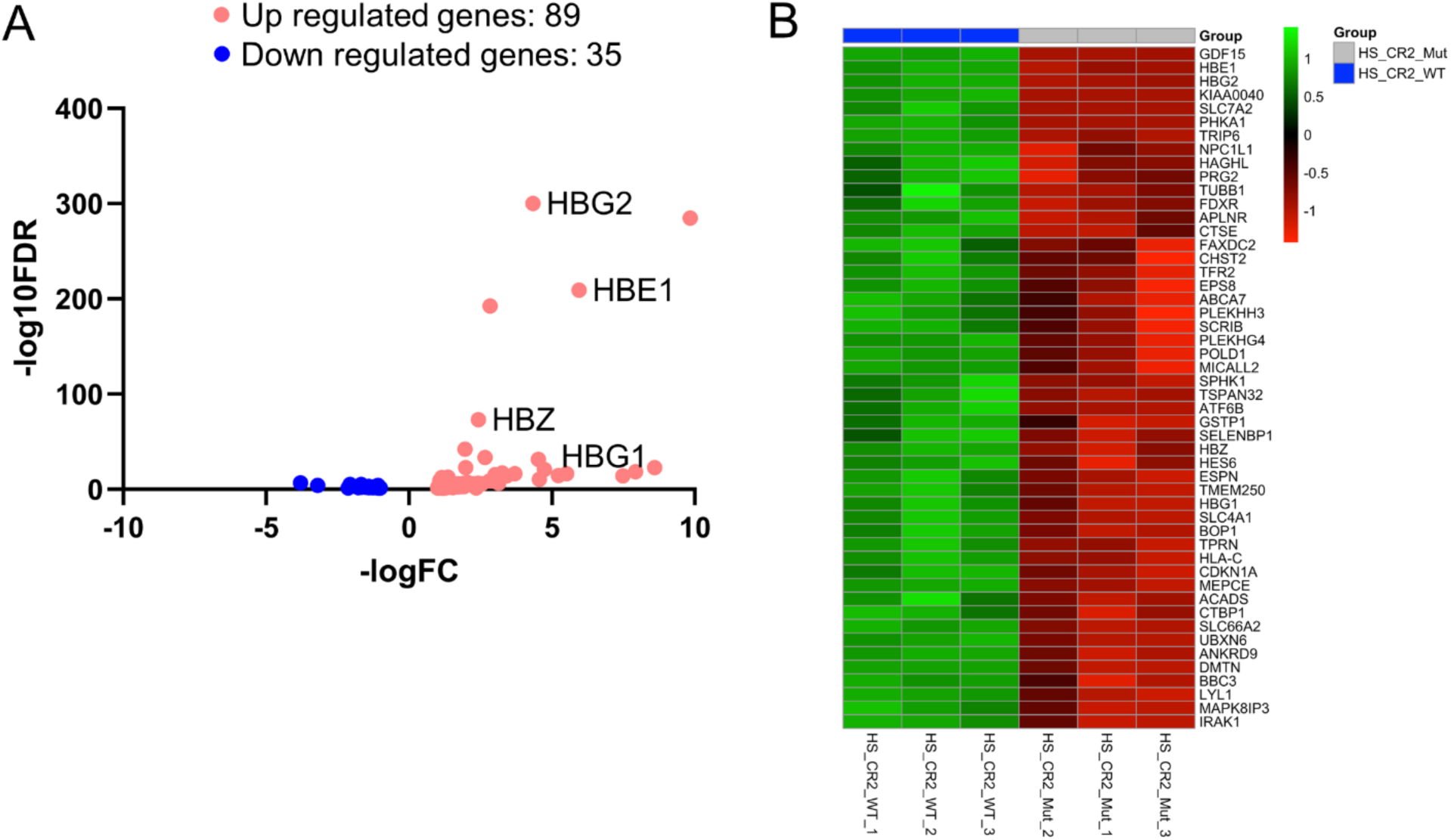
RNA-Seq analysis and CHD4 occupancy of the HBG promoters in CD34+ HSPC derived erythroid expressing wild-type versus mutant CR2 helix peptides. A–B. Volcano plot (A) and heatmap (B) analyses showing comparison of differentially expressed genes (DEGs) in CD34+ HSPC derived erythroid cells expressing wild-type versus mutant CR2 helix peptides.

## Discussion

The central mechanism that enforces silencing of the fetal *HBG* genes and hence HbF expression postnatally is through complexes that occupy the *HBG* promoters and position nucleosomes that block the binding of key activating transcription factors including the NF-Y complex^40^ and GATA1.^41^ The silencing complexes are anchored by the repressive transcription factors BCL11A and ZBTB7A and the MBD2a-NuRD complex which is co-recruited by CpG methylation and the repressive transcription factors that bind there. There are a host of other factors that indirectly affect transcriptional silencing of the *HBG* genes^7^ by either facilitating or inhibiting expression of BCL11A or ZBTB7A, assembly of the silencing complexes or associating with them to potentiate nucleosome positioning.^7,9,42^

Several of the key factors in the central repressor complexes have been targeted to induce therapeutic levels of HbF in the setting of SCD. Gene therapy targeting the occupancy of BCL11A at the *HBG* promoters has been shown to effectively overcome the pathophysiologic manifestations of SCD by inducing high levels of HbF in several clinical trials and is now an approved therapy.^25–27^ Despite the dramatic success of gene therapy, the cost, logistic and sophisticated medical resource issues involved have so far limited its application. It is unlikely that it will be available in the near future for the vast majority of the millions of patients currently affected by SCD worldwide.^43^

This situation has spurred intense research to develop much needed small molecule therapeutics that target the factors that silence HbF expression. While hydroxyurea is clearly beneficial and the best currently available agent, its effects are variable and the levels of HbF it can induce are insufficient to fully stem the morbidity of SCD.^44–46^

With this in mind, we have focused on the MBD2a-NuRD complex as a potential target due to its major role in silencing and pre-clinical data showing that its disruption or depletion of its components results in HbF levels of up to 50% in adult erythroid cells in culture.^13,14^ Structure-function studies have identified critical interaction domains among the MBD2a-NuRD components that could potentially serve as targets for disruption of the complex. We first showed that a coiled-coil interaction between MBD2a and GATAD2A plays a critical role in the formation of NuRD, such that disrupting this interaction induced fetal hemoglobin expression.^12^ However, directly targeting coiled-coil interactions with cell-permeable inhibitors remains a formidable task. Next, we showed that an intrinsically disordered region of MBD2 makes critical interactions with the histone deacetylase core complex of NuRD, and that mutating critical residues at the interface of this interaction induces fetal hemoglobin expression.^47^ Notably, this interaction offers more favorable features for inhibitor development. In the current studies, we continue to focus on the structural components that bridge the histone deacetylase and chromatin remodeling sub-complexes of NuRD by targeting the interaction between the GATAD2A-CR2 domain and the C-terminal region of CHD4.

By employing machine learning structure prediction as a guide, we recently determined the structure of the ternary complex between CHD4, GATAD2A, and CDK2AP1.^37^ These studies revealed that a short central helix in the GATAD2A-CR2 domain binds at the interface between the C1b and C2ab domains of CHD4 and is buried by a short C-terminal helix in CHD4. Hence, we hypothesized that this interaction may be critical for the stability and function of NuRD. Biophysical analyses confirmed that the isolated CR2 helix binds to CHD4 with high affinity and stabilizes the domain as assessed by thermal melting. Furthermore, disrupting helix formation by introducing proline mutations reduces binding affinity and stabilization. Importantly, though, our initial binding analyses involved only the C1bC2a domains of CHD4 since we have not been able to purify constructs that include the C2b domain in isolation. This leaves open the possibility that the interaction with the C-terminal helix of CHD4 could increase the affinity and prohibit competitive inhibition by a peptide or other small molecule. Hence, we tested and found that the isolated CR2 helix does destabilize the ternary complex which includes the CHD4 C-terminal helix. Therefore, the CR2 helix itself can function as a protein-protein interaction inhibitor to disrupt the NuRD complex. Together, the biophysical and structural studies support the possibility of therapeutic targeting of this specific interaction between GATAD2A and CHD4.

As we and others have shown that CHD4 depletion or its dissociation from the core MBD2a-NurD complex can relieve HBG silencing, thus increasing HbF levels,^13,15,29,31,32^ we reasoned that disrupting its interaction with MBD2a-NuRD in HUDEP-2 and adult primary erythroid cells would have similar effects. This was verified by a multipronged approach. Adding back wild-type GATAD2A or a mutant form with amino acid substitutions in the region predicted and validated in structural and biophysical studies showed that mutation of its CR2 helical domain abrogated any silencing of *HBG* in HUDEP-2 adult phenotype erythroid cells. Further, prime editing to introduce the same mutations at L387 and L390 in the endogenous GATAD2A gene in HUDEP-2 cells resulted in dissociation of CHD4 from the core HDCC subcomplex of NuRD, loss of the repressive nucleosomes and repressive histone marks over the *HBG* promoters and high levels of HbF. Likewise, expression of a wild-type CR-2 helical domain peptide, but not a mutant CR-2 peptide bearing the L387/L390P mutations, induced high levels of HbF expression and dissociation of CHD4 from the HDCC subcomplex of NuRD in HUDEP-2 cells. To validate these effects in normal primary adult erythroid cells, the same CR-2 helical peptide was expressed in CD34^+^ HSPCs from multiple origins, followed by erythroid differentiation. This resulted in over an 8-fold increase in HbF levels in erythroid cells expressing wild-type CR-2 peptide compared to those in cells expressing the L387/L390P mutant peptide, with no significant adverse effect on erythroid differentiation.

Among the genetic and epigenetic factors that are essential for maintaining HbF silencing through the central repressive complexes at the *HBG* promoters, most are either transcription factors or require protein-protein interactions for their function. These have traditionally been considered not amenable to small-molecule targeting. The exception is DNA methylation, but despite clinical evidence that hypomethylating agents such as 5-azacytidine are efficacious in SCD,^48,49^ the actual and potential toxicities of these agents have limited their application.

The advancement of PROTAC and other methods for targeting specific proteins for transport to proteosomes for degradation, so-called degrons, has opened a new avenue for the discovery of small-molecule agents that can overcome the limitations on targeting transcription factors.^34^ A molecular glue degron for the epigenetic regulator WIZ has shown preclinical effectiveness and is currently in clinical trials,^16^ as is a degron targeting both WIZ and ZBTB7A/LRF. Preclinical studies aimed at targeted degradation of the major silencing transcription factor BCL11A have been reported.^50,51^

An alternative approach to disrupting the central silencing complexes at the *HBG* promoters is to develop a therapeutic interfering peptide to dissociate CHD4 from the MBD2a HDCC NuRD subcomplex, as supported by the on-target proof-of-principle findings in this report. Delivery of natural peptides is limited by issues of the requirement for parenteral administration and difficulties in intracellular delivery without proteasomal degradation. These limitations can be overcome by advances in the synthesis of macrocyclic small peptides containing non-natural amino acids and moieties that facilitate cell permeability.^52–56^ There is already a growing number of approved and clinically effective macrocyclic peptides, some of which are orally administered. Thus, such a peptide based on the biologically active CR-2 helical domain peptide disrupter shown here to disrupt the interaction to induce high levels of HbF in primary human erythroid cells could be an effective therapeutic agent in SCD. This approach is being pursued by optimizing the macrocyclic peptide that was shown to disrupt the ternary complex between GATAD2A and CHD4 in our recent related studies.^37^

Since any of the factors that constitute the central *HBG* silencing complex or those that affect their action are likely to be involved in other biological processes, there is the potential for unintended off-target toxicities of agents that inhibit, degrade, or disrupt them. Given the relatively mild phenotype of MBD2 knockout mice,^28^ and the high level of Hb F induced by disruption of the MBD2a-NuRD complex, it may represent a successful target for therapeutic induction of HbF expression in SCD. The ideal small-molecule agent for the treatment of SCD will be determined empirically by studies that demonstrate which agent(s) balance high-level HbF induction with minimal toxicity. This reality encourages further exploration of multiple HbF silencing factors as potential therapeutic targets. The results reported here showing the ability of a small peptide from the CR2 domain of GATAD2A to induce high levels of HbF in primary adult erythroid cells without perturbing differentiation support the pursuit of this particular approach.

## Acknowledgements

This work was supported by grants from the National Institutes of Health award R01 DK115563 to DCW and GDG. Core facilities were supported by the National Cancer Institute of the National Institutes of Health under award number P30 CA016086 to the UNC Lineberger Comprehensive Cancer Center and under award number P30 CA16059 to the VCU Massey Comprehensive Cancer Center. The authors acknowledge additional generous support provided by Eshelman Innovation at the UNC Eshelman School of Pharmacy, the UNC Lineberger Comprehensive Cancer Center (LCCC) and the VCU Massey Comprehensive Cancer Center (MCCC).

## Author Contributions

GDG, DCW and SS designed experiments, SS, GD-D, SG, TL, HW, CT and DCW carried out experiments, SS, DCW, GG, and MD analyzed data and SS, GDG and DCW wrote the manuscript.

## Disclosure of Conflicts of Interest

The authors declare no conflicts of interest.

## Notes

### Competing Interest Statement

The authors have declared no competing interest.

