## Supplemental Table 1 for "Structure-guided targeting of the GATAD2A-CHD4 interaction within the MBD2-NuRD complex results in high levels of HbF in adult erythroid cells"

### Supplement figure legend

**Supplement Figure 1. Prime editing of GATAD2A at L387 and L390 site.**

A. Sanger sequencing result of PE edited homozygous HUDEP-2 clone. PE-CR2 DLP is the homozygous editing of GATAD2A L387P and L390P.

B. Q-PCR results of differentiation marker CD71 and CD235 between PE editor control and G2A mutant.

C. qPCR results showing the γ/(γ+β)% mRNA ratio and the relative expression levels of globin mRNAs in HUDEP-2 cells and Prime Editor (PE)-edited single-cell clones. "PE-G2A bulk" refers to the bulk population edited by Prime Editing. "No edit" represents the wild-type control. "L387 or L390 only" is single clones with a single amino acid substitution. "Heter L387390P" is a heterozygous mutant with substitutions at both L387 and L390. The left panel shows the γ-globin mRNA ratio; the middle and right panels show the relative expression levels of γ-globin and β-globin mRNAs, respectively. PE editing led to a modest increase in γ-globin expression, while β-globin levels remained unchanged or were not decreased. Statistical analysis of the violin plot data was performed using the Kruskal–Wallis one-way ANOVA test. *P<0.05, ** P<0.01, *** P<0.001.

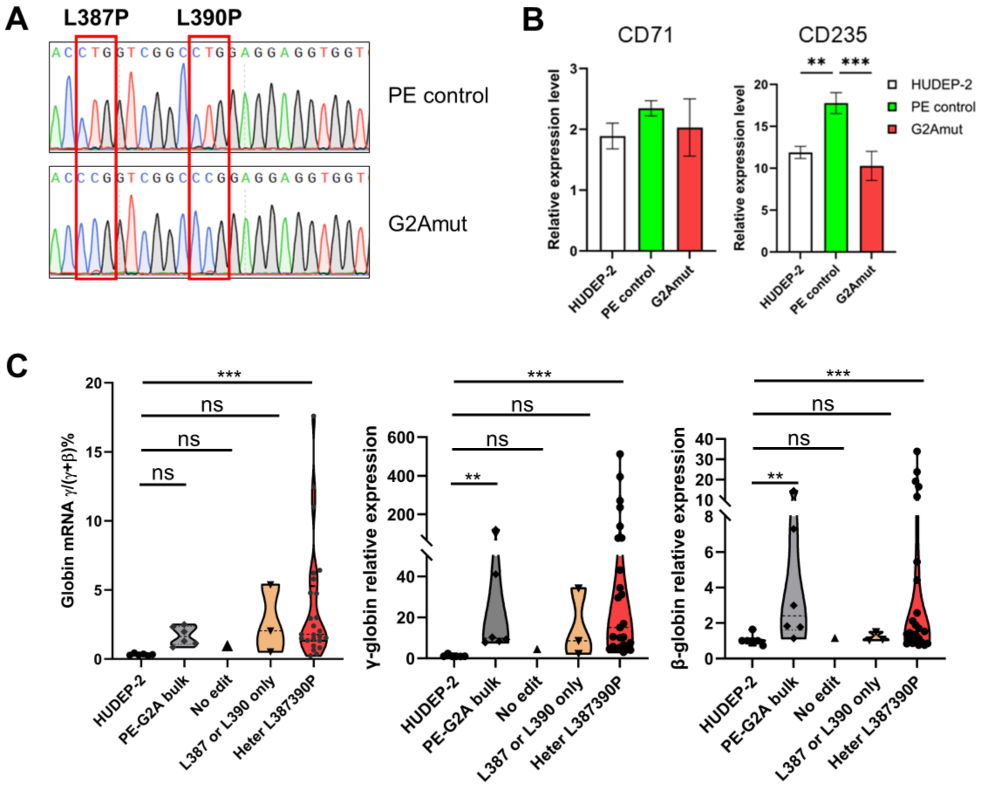
